# The dual role of N6-methyladenosine on mouse maternal RNAs and 2-cell specific RNAs revealed by ULI-MeRIP sequencing

**DOI:** 10.1101/2021.12.13.472368

**Authors:** You Wu, Xiaocui Xu, Meijie Qi, Chuan Chen, Mengying Li, Rushuang Yan, Xiaochen Kou, Yanhong Zhao, Wenqiang Liu, Yanhe Li, Xuelian Liu, Meiling Zhang, Chengqi Yi, Hong Wang, Bin Shen, Yawei Gao, Shaorong Gao

## Abstract

N^6^-methyladenosine (m^6^A) and its regulatory components play critical roles in various developmental processes in mammals(*1-5*). However, the landscape and function of m^6^A in the maternal-to-zygotic transition (MZT) remain unclear due to limited materials. Here, by developing an ultralow-input MeRIP-seq method, we revealed the dynamics of the m^6^A RNA methylome during the MZT process in mice. We found that more than 1/3 maternal decay and 2/3 zygotic mRNAs were modified by m^6^A. Moreover, m^6^As are highly enriched in the RNA of transposable elements MTA and MERVL, which are highly expressed in oocytes and 2-cell embryos, respectively. Notably, maternal depletion of *Kiaa1429*, a component of the m^6^A methyltransferase complex, leads to a reduced abundance of m^6^A-marked maternal RNAs, including both genes and MTA, in GV oocytes, indicating m^6^A-dependent regulation of RNA stability in oocytes. Interestingly, when the writers were depleted, some m^6^A-marked 2-cell specific RNAs, including *Zscan4* and MERVL, appeared normal at the 2-cell stage but failed to be decayed at later stages, suggesting that m^6^A regulates the clearance of these transcripts. Together, our study uncovered that m^6^As function in context-specific manners during MZT, which ensures the transcriptome stability of oocytes and regulates the stage specificity of zygotic transcripts after fertilization.

**One Sentence Summary:** m^6^A RNA methylation stabilizes the maternal RNAs in mouse oocytes and degrades the 2-cell specific RNAs in the cleavage-stage embryos.

## Main Text

Fertilization triggers a remarkable and complex cell fate transition from oocytes to totipotent embryos, termed the maternal-to-zygotic transition, including dramatic remodeling of the chromatin epigenome, transcriptome and proteome(*6*). However, in mice, transcription is effectively silenced from fully grown germinal vesicle (GV) oocytes until ZGA at the late 1-cell stage. Therefore, post-transcriptional modifications regulate the storage, timely decay and sequential activation of maternal transcripts, which ensures the switch from maternal to embryonic control of gene expression.(*7, 8*) Although some reprogramming details of the epigenetic and chromatin landscape have been described in recent years, the various control mechanisms on RNA remain elusive.

In eukaryotes, N^6^-methyladenosine (m^6^A) is found on messenger RNAs (mRNAs), repeat RNAs and long noncoding RNAs (lncRNAs)(*3, 4, 9-11*) and participates in various important biological events by playing roles in RNA-related processes and chromatin state regulation(*1, 2, 5, 12-15*). A previous study in *zebrafish* found that m^6^A could promote maternal RNA degradation during the MZT process(*16*). In mammals, depletion of m^6^A reader proteins, such as Ythdc1 and Ythdf2, or deficiency of m^6^A writers, such as Mettl3/14/16 and Kiaa1429, leads to developmental defects in oocytes or early embryonic lethality(*17-23*), suggesting that m^6^A may play important roles in oocyte and early embryo development. However, genome-wide profiling and the potential roles of m^6^A in this process have been largely untacked due to the limited number of oocytes and preimplantation embryos. In this study, we first developed an ultralow-input MeRIP-seq (ULI-MeRIP-seq) method, which enabled the profiling of m^6^A RNA methylation with 50 ng of total RNA. With this method, we revealed the dynamics of the m^6^A landscape of the transcriptome in the MZT process from GV oocytes to 4-cell stage embryos, which covered oocyte maturation, fertilization and ZGA events in the MZT process.

### RNA m^6^A methylome in mice MZT

To investigate the role of RNA m^6^A methylation during MZT in mammals, we optimized the protocol of m^6^A RNA immunoprecipitation followed by high-throughput sequencing (MeRIP-seq)(*24*) and developed ultralow-input MeRIP-seq (ULI-MeRIP-seq), which can examine the m^6^A RNA methylome using 50 ng of total RNA (Fig. 1A and fig. S1A, see methods for details). In the preliminary experiments, 50 ng and 2 µg of total RNA from mouse ES cells was used for MeRIP-seq, and the IP efficiency was validated by qPCR of GLuc/CLuc spikes-in and endogenous mRNA *Klf4(19*) (fig. S1, B and C). The modification on pluripotency marker genes was also consistent with the previous reports(*19, 25*) (fig. S1D). The MeRIP-seq data generated using 50 ng total RNA were high-quality and recapitulated the results generated using 2 µg total RNA (fig. S1, E to I). We then extracted RNA samples in the MZT process, including oocytes at the GV and metaphase II (MII) stages, as well as embryos at the late 1-cell (L1C), late 2-cell (L2C) and 4-cell (4C) stages and performed LS-MS/MS (Fig. 1B). The overall abundance of m^6^A continuously decreased during MZT, which was quantified by LC-MS/MS (fig. S2A). We then employed ULI-MeRIP-seq to profile m^6^A with 50 ng total RNA for each stage with two or three replicates, and total RNA-seq of relevant samples without IP reaction was used as input. PCA of genes showed that both the transcriptome and m^6^A methylome diverged after 2-cell stage when major ZGA occurred (Fig. 1C). The ULI-MeRIP-seq data showed highly efficient enrichment for m^6^A, and all replicates, including input libraries, were highly comparable (Fig. 1C, fig. S2, B to D, and tables S1 and S2). Saturation analysis showed that our data were able to detect both high-level and low-level transcripts of genes (fig. S2E). To identify m^6^A-enriched sites, we performed peak calling using model-based analysis (MACS). As expected, the length distribution and enrichment of canonical [RRACH] motif (R= G/A; H= A/C/U) on m^6^A peaks at each stage was consistent with previous studies(*9, 10*); however, the density of [RRACH] at exon peaks decreased over the MZT process (fig. S3, A to C). In addition to the well-characterized enrichment of m^6^A methylation at coding sequences (CDSs), untranslated regions (UTRs), and near stop codons (Fig. 1D), we found that a considerable number of m^6^A peaks were mapped to distal intergenic regions, which was particularly high at the late 1-cell stage (fig. S3D).

**Fig. 1.**
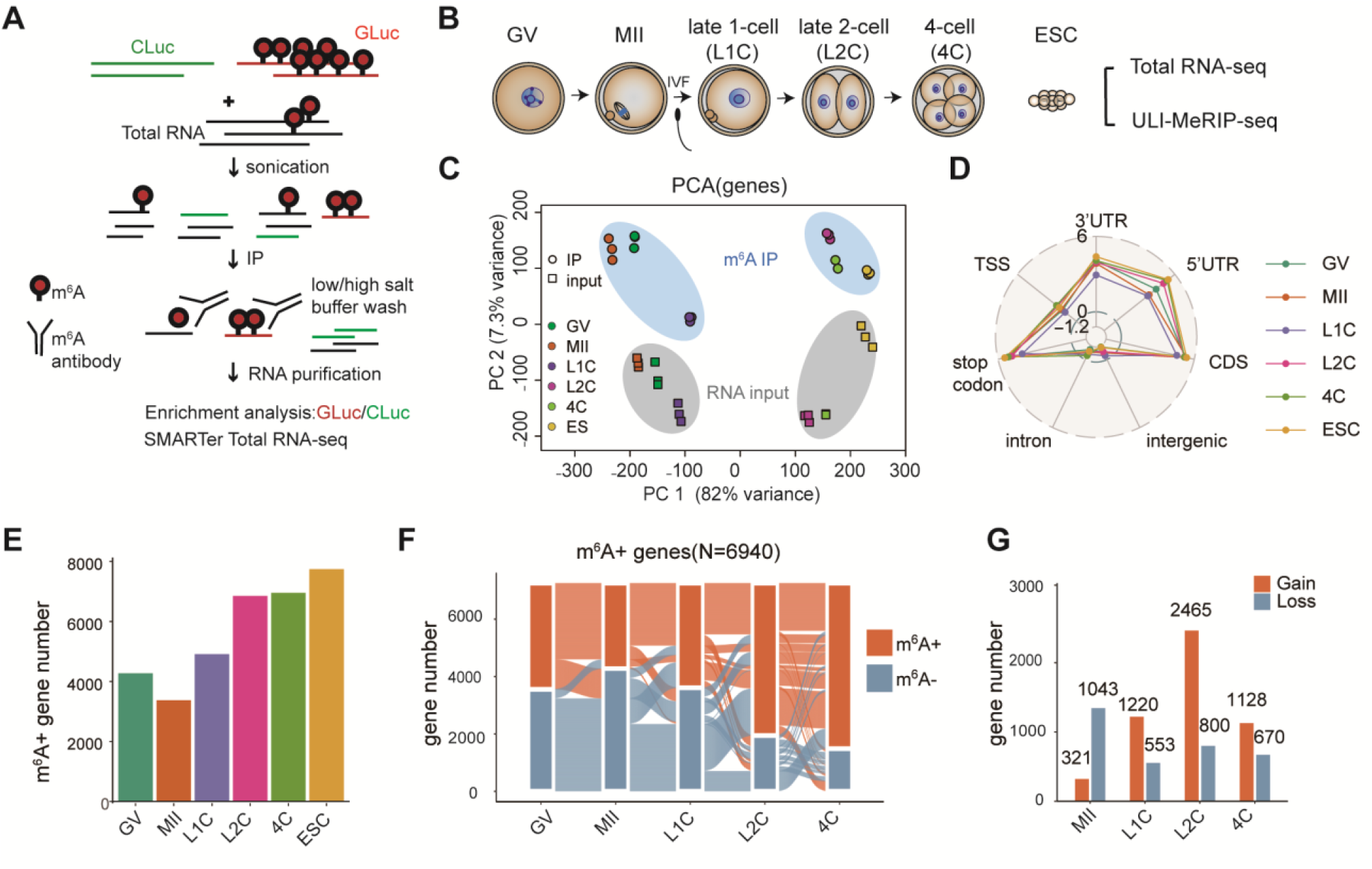
RNA m^6^A modification in mouse oocytes and early embryos. **(A)** Schematic diagram of the ULI-MeRIP-seq procedure. Spike-in RNAs, including unmodified control RNA (Cypridina Luciferase, CLuc) and m^6^A methylated control RNA (Gaussia luciferase, GLuc), were added to total RNA to monitor m^6^A enrichment (see methods). **(B)** GV and Mil oocytes, late 1-cell, late 2-cell, 4-cell and mESCs were collected for total RNA-seq and ULI-MeRIP-seq. **(C)** Principal component analysis (PCA) of m^6^A and input RNA data of genes for all samples. **(D)** Radar chart showing the enrichment score of m^6^A peaks at TSS, 3’UTR, 5’ UTR, CDS, intergenic, intron and stop codon. **(E)** The number of genes harboring m^6^A (m^6^A+) in each stage. **(F)** Alluvial diagram showing the global dynamics of m^6^A+ genes during MZT (N=6940). Each line represents a gene that is classified as a m6A+ class in at least one stage. **(G)** The number of genes gaining or losing m^6^A at each stage compared to the prior stage during MZT. The number of genes gaining or losing m^6^A is indicated above each bar.

We then defined the m^6^A-marked (m^6^A+) genes at each stage and calculated the dynamics of m^6^A peaks and m^6^A+ genes during the MZT process. To our surprise, compared to the continuously decreased m^6^A/A ratio detected by LS-MS/MS, m^6^A peaks as well as m^6^A+ genes increased after fertilization (Fig. 1, E to G and fig. S3, E to G), suggesting that de novo m^6^A modification can be established along with ZGA. Interestingly, total m^6^A peaks doubled shortly after fertilization and possessed more striking dynamic changes in the 2-cell and 4-cell stages compared to those of m^6^A+ genes. Detailed analysis revealed that the ratio of m^6^A peaks on transposons significantly increased in the gain peaks at late 1-cell and lost peaks at 2-/4-cell stages (fig. S3H). These data reveal a highly dynamic landscape of the m^6^A methylome during MZT in which de novo establishment and severe loss of m^6^A may occur on both genes and transposon RNAs.

### m^6^A marks maternal decay and ZGA genes in MZT

To investigate the association between m^6^A methylation and the MZT process, we first analyzed the transcriptome data and identified 2,533 maternal decay genes (termed MD genes) and 1,115 ZGA genes based on their abundance change over the MZT process (Fig. 2A, see methods and table S3). We then found that 43% of MD genes were marked by m^6^A in GV or MII oocytes, which exhibited a decreased m^6^A signal around the stop codon throughout the MZT process (Fig. 2A and fig. S4A). Gene ontology (GO) analyses showed that m^6^A unmarked (m^6^A -) MD genes were significantly enriched in energy metabolism pathways, including mitochondrial organization and cellular respiration, such as the Nduf family and Atp family (Fig. 2B and fig. S4B). Meanwhile, m^6^A+ MD genes were mainly associated with TGF-β, epithelial to mesenchymal transition and GTPase signaling pathways, including *Gdf9, Tgfb2, Jun, Mos* and *Bmp15*, which play essential roles in oogenesis (Fig. 2, B and C, and fig. S4B). More detailed analysis revealed that the m^6^A+ MD genes possess relatively higher [RRACH] density, and their abundance was higher in oocyte stages, which was consistent with the observation in *zebrafish*(*16*) (Fig. 2D and fig. S4D).

**Fig. 2.**
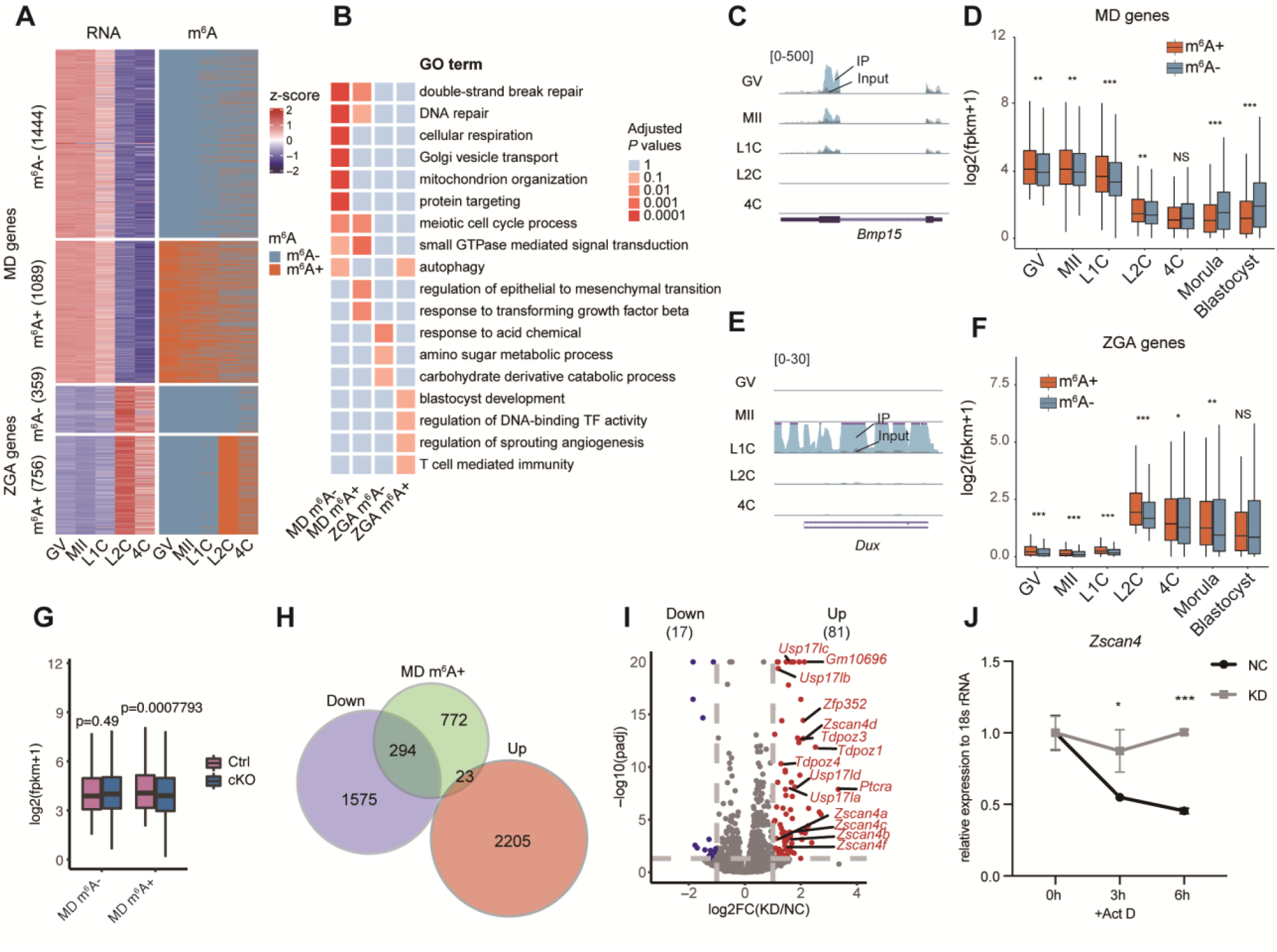
Association between m^6^A dynamics and maternal decay genes and ZGA genes during MZT. **(A)** Heatmap showing normalized gene expression (left panel) and m^6^A modification status (right panel) of maternal decay (MD) genes and ZGA genes across different stages. 0 and 1 represent genes without or with m6A peaks, respectively. **(B)** Gene ontology (GO) analysis of the 4 groups of genes according to Fig 2A. GO terms for each functional cluster were summarized to a representative term, and adjusted p-values were plotted lo show the significance **(C)** The UCSC browser track showing IP and input reads of m^6^A+ MD gene Bmp15. **(D)** Boxplot showing the expression levels of m^6^A+ and m^6^A-MD genes during early embryo development **(E)** The UCSC browser track showing IP and input reads of m^6^A-+ ZGA gene Dux. **(F)** Box plot showing the expression levels of m^6^A-+ and m^6^A-ZGA genes during early embryo development **(G)** Box plot showing the expression levels of m^6^A-end m^6^A+ MD genes in control and *Kiaa1429* cKO GVs. **(H)** Venn plot showing the overlapping gene number or DEGs in *Kiaa1429* cKO GV and maternal m^6^A+ MD genes. **(I)** Volcano plot showing DEGs in morula after m^6^A writer Mettl3, Mettl14 and Mettl16 knockdown Names of representative 2C genes are marked. KD, writer knockdown: NC, nontarget **(J)** Decay rate of *Zscan4* RNA at 4-cell stage. qPCR showed the RNA level of *Zscan4* relative to 18S rRNA at the 4-cell stage with Actinomycin D (Act D) treatment in nontarget and writer knockdown embryos. Error bars indicate mean ± SD.

Interestingly, we found that approximately 68% of ZGA genes were modified by m^6^A in the late 1-or 2-cell stages, which indicates that m^6^A was installed along with the transcription of ZGA genes (Fig. 2A and fig. S4A). Compared to the m^6^A-ZGA genes, which are predominately enriched for acid chemical and amino sugar metabolic process like *Npl* (fig. S4C), m^6^A+ ZGA genes were strongly enriched for processes involved in blastocyst development and transcription factor activity, such as Tet1 (Fig. 2B). Notably, 2-cell marker genes, including *Dux* and *Zscan4*, were found to be marked by m^6^As when they were reactivated in minor ZGA (Fig. 2E and fig. S4C). Additionally, the [RRACH] density on m^6^A+ ZGA genes was higher, and the ZGA genes with m^6^A possessed higher levels of RNA in early cleavage embryos (Fig. 2F and fig. S4D).

### Kiaa1429-mediated m^6^A modification stabilizes maternal mRNAs

The methylation of m^6^A is mainly dependent on the methyltransferase complex, in which METTL3 and METTL14 work as catalytic cores and interact with WTAP, KIAA1429 (also known as VIRMA), RBM15/RBM15B, ZC3H13 and other undefined proteins(*3, 26*). A previous study showed that KIAA1429 is essential for m^6^A deposition near stop codons and 3’UTRs in human cells (*27, 28*). Our collaborators found that loss of Kiaa1429 in oocytes could lead to abnormal RNA metabolism in GV oocytes and abolished the ability to undergo germinal vesicle breakdown (GVBD)(*21*), suggesting that Kiaa1429-mediated RNA metabolism was important for maternal RNAs in mice. To understand the mechanism, we performed RNA-seq and ULI-MeRIP-seq of GV oocytes from *Kiaa1429*^*Zp3*^ control and cKO mice (fig. S5, A and B). Consistent with a previous study(*21*), both the transcriptome and m^6^A methylome were impacted, and the m^6^A modification in GV oocytes decreased dramatically when Kiaa1429 was deleted (fig. S5, B to D). Further analysis revealed that the RNA abundance of m^6^A-marked MD genes was reduced significantly in Kiaa1429-deficient oocytes compared to unmarked genes (Fig. 2G), suggesting a potential correlation between m^6^A and high abundance of RNA transcripts in oocytes, which was observed in maternal decay genes. We then identified the differentially expressed genes (DEGs) between control and *Kiaa1429* ^*Zp3*^ cKO oocytes, in which 1869 genes were repressed and 2228 genes were upregulated upon Kiaa1429 depletion (Fig. 2H, fig. S5E, and table S4). We found that 92.7% of m^6^A-marked MD DEGs were downregulated (294/317), suggesting the role of m^6^A in maintaining high levels of those genes (Fig. 2H). To verify the change in transcriptional activity on m^6^A-marked genes, we performed ATAC-seq with the nuclei of control and *Kiaa1429*^*Zp3*^ cKO GV oocytes. The nucleus of non-surrounded nucleolus (NSN) and partly surrounded nucleolus (PSN) GV were collected and compared respectively. We compared chromatin accessibility at the promoter region (3 kb around the TSS) of m^6^A+ MD genes and found that the profiles of ATAC-seq signals between control and *Kiaa1429*^*Zp3*^ cKO oocytes were almost comparable, suggesting that the transcriptional activity of m^6^A-marked genes was not apparently affected (fig. S5, F and G). Therefore, the reduced RNA transcripts on m^6^A-marked genes may come from the reduced stability of RNA but not caused by the regulation of transcription activity. All these data demonstrate that Kiaa1429 is essential for m^6^A deposition in GV oocytes and subsequent stabilization of m^6^A-marked maternal transcripts.

Next, we wondered whether Ythdf2, like in *zebrafish*, is involved in the RNA decay of maternal RNAs after fertilization in mice. We used previously established *Ythdf2*^*Zp3*^ cKO mice and observed 2-cell embryo arrest, which is similar to the published study(*17*). Single-cell RNA-seq was performed using MII oocytes and 2-cell embryos from *Ythdf2*^*Zp3*^ control and cKO mice (fig. S6, A and B). The heatmap showed that Ythdf2 depletion in oocytes repressed the degradation of maternal decay genes in the 2-cell stage but did not impact RNA abundance in MII oocytes (fig. S6C). However, to our surprise, not only the degradation of m^6^A+ but also m^6^A-maternal decay genes appeared to be impacted by Ythdf2 depletion (fig. S6D). Therefore, the function of Ythdf2 in MZT may not be limited to its direct binding to m^6^A but in a more complex manner.

### m^6^A regulates the decay of 2-cell specific ZGA genes

In *zebrafish*, m^6^A was also observed on ZGA genes(*16*); however, compared to maternal decay genes, the roles of m^6^As on zygotic mRNAs were barely discussed. As the removal of m^6^A in GV oocytes can lead to severe defects in folliculogenesis, we have to establish a new platform to realize the inhibition of de novo m^6^A installation on ZGA genes. Here, we analyzed the expression pattern of the reported m^6^A methyltransferases and found that *Mettl3* and *Mettl14* were highly expressed in MII oocytes, but all these genes, including *Mettl16*, possessed considerable RNA levels in the late 1-cell stage (fig. S7A). Therefore, we injected siRNA or antibodies against all three Mettl genes into MII oocytes to inhibit the function of these proteins (fig. S7B). As a result, the RNA level of *Mettl3/14/16* was reduced significantly, and the development rate decreased modestly (fig. S7, C and D). RNA-seq revealed that the RNA level of m^6^A+ ZGA genes in KD embryos was unchanged in the 2-cell stage but increased at the morula stage (fig. S7E). Transcriptome-wide analysis showed that the transcriptome of KD embryos was almost comparable to that of control embryos at the 2-cell stage, but in the morula stage, a group of genes were significantly upregulated upon KD of m^6^A methyltransferases (Fig. 2I, fig. S7F, and table S5). Interestingly, most of the genes upregulated in KD morula were 2-cell specific ZGA genes with m^6^A, and many 2C marker genes, including the Zscan4 family, were included and possessed higher RNA levels in KD morula (Fig 2i and Extended Data Fig 7g-h). With these data, we hypothesized that m^6^A may direct the RNA decay of 2-cell specific ZGA genes to ensure the stage-specific expression of these genes. To prove this, we first confirmed the decay of Zscan4 RNA in control embryos at the 4-cell stage when Actinomycin D (1 µg/ml or 5 µg/ml) was used to inhibit transcription (fig. S7I). We then verified that the decay of Zscan4 RNA was impaired when the function of m^6^A methyltransferases was inhibited (Fig. 2J). In summary, our data indicate that installation of m^6^A on 2-cell specific ZGA genes promoted the clearance of their transcripts to ensure their stage specificity in embryos.

### Enrichment of m^6^A on the transcripts of transposable elements

Transposable elements (TEs), mainly retrotransposons, are an important component and occupy more than one-third of the mouse genome(*29*). The majority of TEs are long interspersed nuclear elements (LINEs) or short interspersed nuclear elements (SINEs), and approximately 10% of them are long terminal repeat (LTR) elements, which resemble retroviruses. Many studies have reported the dynamic stage-specific expression of certain retrotransposons in mouse germ cells and preimplantation embryos, and some of them, such as MaLR, MERVL and LINE1, reflect a widespread mechanism for regulating MZT and embryo development(*6, 30, 31*). Recently, m^6^A was found to be enriched on the RNA of retrotransposons and regulates their half-life or their interaction with proteins(*12-15*). In our data, we noticed the existence and dynamics of m^6^A peaks on transposons in the MZT process but called for systematic analysis.

To start this part, we first isolated the reads located on repetitive RNA and calculated the enrichment of m^6^A on the subfamilies of TEs (see Methods and fig. S8A). We then calculated the m^6^A enrichment on different classes of repeats. Generally, m^6^A methylation was mainly enriched in LTR RNAs in all MZT samples. However, m^6^As in ESCs was mainly enriched in LINE RNAs (fig. S8B). Subsequently, we analyzed the m^6^A enrichment and expression of the main family in LTR and LINE. As shown, MaLR and ERVL, LTR class III retrotransposons, possess highly enriched m^6^As in oocytes and early cleavage embryos, respectively (fig. S8C), which is consistent with their stage-specific expression features (fig. S8D) (*30*).

To be more precise, we defined all the expressed repeat subfamilies in the MZT process (expression level >5) (Fig. 3A) and compared their expression and m^6^A dynamics. These TE families were classified into three clusters, termed maternal TEs, ZGA TEs and mid-preimplantation gene activation (MGA) TEs, based on their expression pattern from GV oocytes to blastocyst embryos (Fig. 3A). Most of the maternal TEs, such as MTAs (MTA_Mm and MTA_Mm_int), MT-int and RLTRs, possess high RNA abundance and enrichment of m^6^A from oocytes to late 1-cell embryos (Fig. 3A). Meanwhile, all the ZGA TEs, including MERVL (MERVL_int and MT2_Mm), MT-int and ORR1As, possessed high RNA abundance and enrichment of m^6^A from late 1-cell to 4-cell embryos (Fig. 3A). This observation was similar to the m^6^A+ MD and ZGA genes. Therefore, the role of m^6^A on retrotransposon RNAs might be similar to that we discovered on coding mRNAs.

**Fig. 3.**
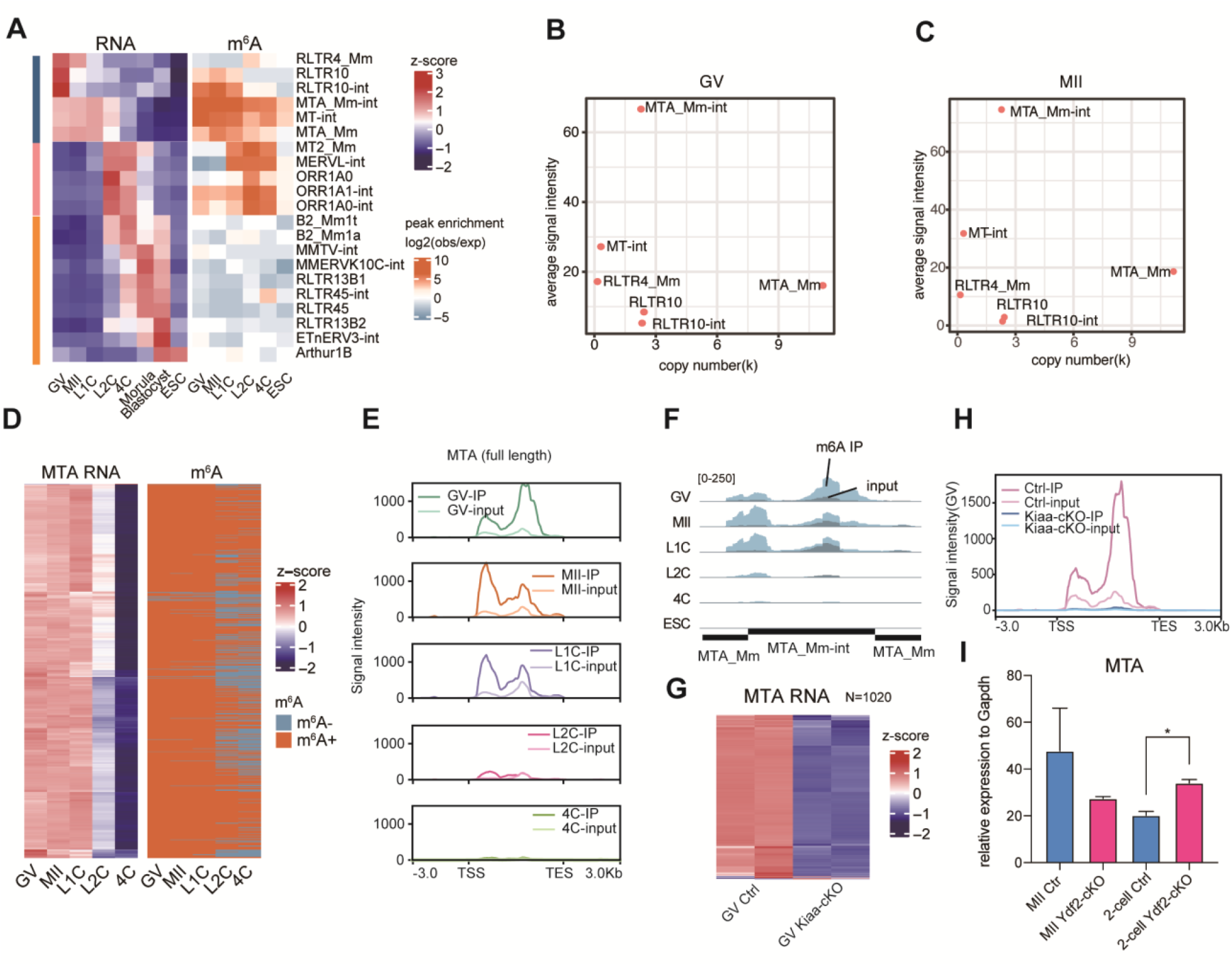
M^6^A is indispensable for the high RNA level of maternal TE MTA. **(A)** Heatmap of normalized RNA levels (left panel) and m^6^A enrichment scores (right panel) of all expressed transposon elements. Expressed transposon elements were defined as those with a mean expression level greater than 5. Pink, yellow and dark blue bars on the left represent 3 clusters of TEs according to expression pattern. (**B** and **C**) Plots showing average RNA level and genomic copy number for maternally expressed TEs in GV and Mil, respectively. **(D)** Heatmap showing the normalized RNA level and m^6^A peak status of full-length MTA copies. **(E)** Average profile of m^6^A IP and input signal of full-length MTA during MZT. **(F)** The UCSC browser track showing IP and input reads of MTA during MZT. **(G)** Heatmap showing the normalized MTA RNA level of each copy during MZT in WT mice and GV oocytes of *Kiaa1423* control and maternal cKO mice. **(H)** Average m^6^A IP and input signal in control and *Kiaa1429* Zp3-cKO GV oocytes at MTA. The IP and input signal were scaled by GLuc and CLuc spike-ins, respectively. **(I)** MTA RNA levels in Mil oocytes and 2-cell embryos of control and *Ythdf2* maternal cKO mice quantified by qPCR. Error bars indicate mean ± s.e.m.

### m^6^A facilitates the storage and decay of MTA mRNAs

Among the subfamilies of maternal TEs, MTA, phylogenetically the youngest and most abundant mouse transcript (MT) subfamily, exhibited the highest RNA intensity in oocytes as well as its abundance in the genome (Fig. 3, B and C). Previous studies revealed oocyte-abundant mouse transcript (MT) transposable elements, which are involved in the MaLR family, accounted for over 12% of the total ESTs in the GV oocyte cDNA library(*30*). Moreover, knockdown of MT transcripts in GV oocytes or zygotes with pronuclei (PN) affects the GVBD rate or causes cleavage arrest(*32*), suggesting that MT transposable elements play critical roles in oocyte maturation and early embryo development. Therefore, we mainly focused on MTA in further study.

By checking individual TE copies, we testified that almost all intact MTAs with high expression, divided into 5 clusters based on the expression level, are marked by m^6^A peaks from GV oocytes to late 1-cell embryos, in which canonical [RRACH] motifs were highly enriched (Fig. 3D and fig. S8, E and F). The profiling of m^6^A on MTA during MZT not only showed continuous demethylation after the zygote stage but also exhibited a significant m^6^A density shift from TES to TSS in MII oocytes and late 1-cell embryos (Fig. 3, E and F). Moreover, this result can be repeated when unique mapping was performed for data analysis (data not shown). Considering that the MTA RNAs stored in GV oocytes will readily decay in matured or fertilized oocytes, the “shift” of m^6^A peaks suggests that region-specific m^6^A deposition might be important for m^6^A-mediated regulation.

We also checked the behavior of MTA transcripts in *Kiaa1429*^*Zp3*^ cKO oocytes. In addition to the loss of m^6^A, the dramatically decreased RNA abundance on MTA copies was much more severe than that observed on m^6^A-marked genes when Kiaa1429 was depleted (Fig. 3, G and H and fig. S8G). We also compared the ATAC-seq signals of the clustered MTAs with different expression levels and excluded the potential impact of transcriptional activity (fig. S8H). These results indicate that m^6^As on MTA elements was extremely important for maintaining a high abundance of MTA RNA in GV oocytes, and the loss of MTA RNA may also contribute to the abolished oocyte competence in Kiaa1429 cKO mice.

We further checked the RNA level of MTA in MII and 2-cell samples and found that the decay of MTA was blocked in *Ythdf2*^*Zp3*^ cKO embryos, which suggests that Ythdf2 regulates the degradation of MTA in fertilized embryos (Fig. 3I).

### m^6^A regulates the decay of MERVL RNA

Among the subfamilies of defined ZGA TEs, MERVL, the most representative 2-cell retrotransposon element, exhibited the highest RNA intensity and genome abundance in 2-cell and 4-cell stage embryos (Fig 4a-b). Again, we divided all intact MERVL copies into 5 clusters based on their expression level and calculated the m^6^A profile on MERVL. Here, we observed that the deposition of m^6^A appeared at the late 1-cell stage and increased in the 2-cell and 4-cell stages (Fig. 4C and fig. S9, A to D). Similar to the 2-cell marker gene *Zscan4*, the RNA level of MERVL at the morula stage also increased significantly when the function of m^6^A methyltransferases was impaired (Fig. 4D and fig. S9E). A decay assay at the 4-cell stage demonstrated that RNA decay of the MERVL transcripts was inhibited when m^6^A writers were knocked down (Fig. 4E). To further validate the direct regulatory role of m^6^A on MERVL RNAs, we employed the dCas13b-ALKBH5 system in embryos(*33*) (fig. S9F). By coinjection of dCas13b-ALKBH5 mRNA and multiple gRNAs targeting MERVL (fig. S9G), we observed decreased m^6^A on MERVL at the 2-cell stage embryos and increased MERVL transcripts at the 4-cell stage, which was validated by MeRIP-qPCR and RT-qPCR, respectively (Fig. 4F and fig. S9H). Our data uncovered a conservative regulatory mechanism on ZGA genes and TEs, in which m^6^A regulated the decay of MERVL in 4-cell stage embryos.

**Fig. 4.**
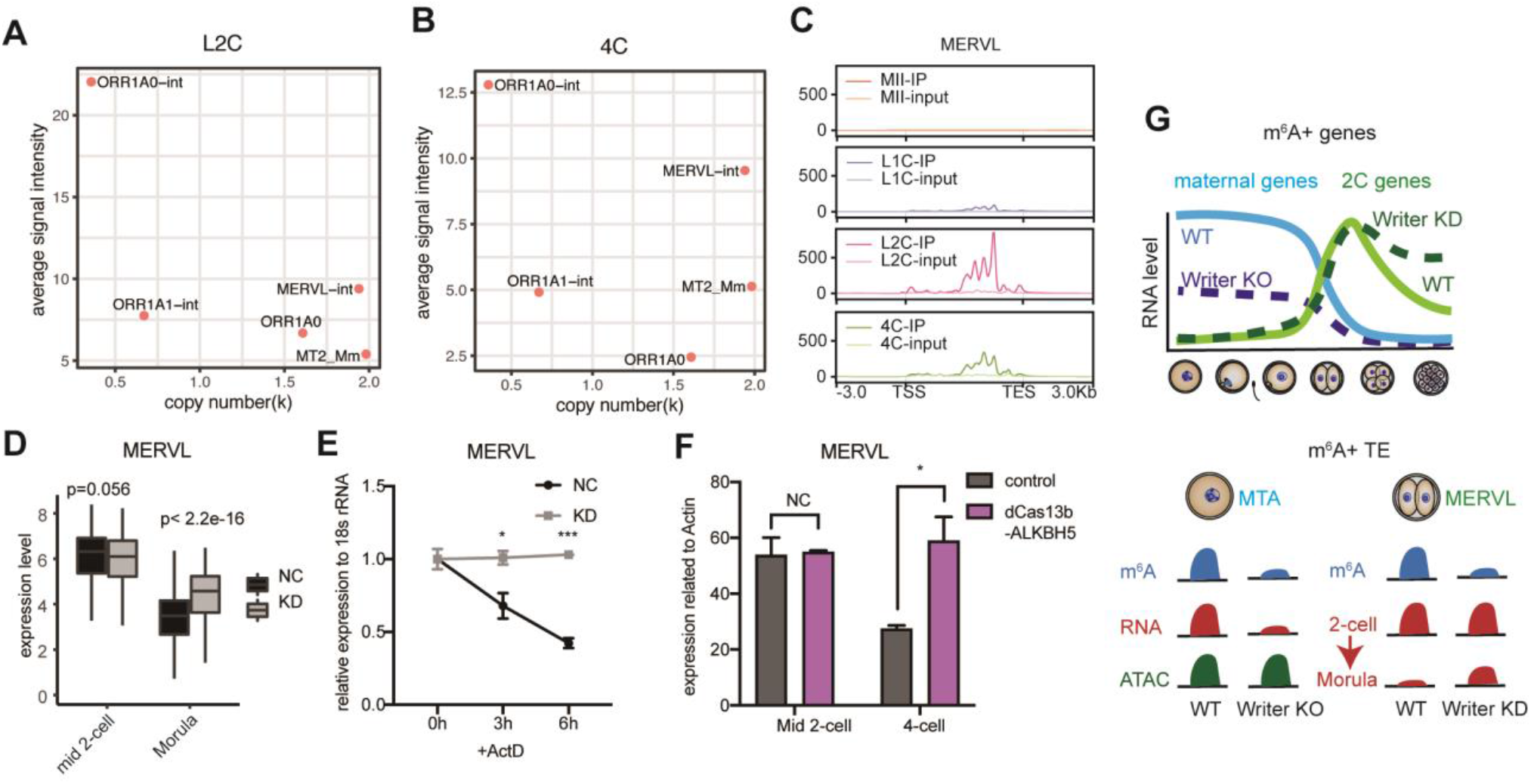
M^6^A promotes degradation of MERVL transcripts. (**A** and **B**) Plots showing the average RNA level and genomic copy number for ZGA TEs in 2-cell and 4-cell embryos, respectively. **(C)** Average profile of m^6^A IP and input signal of ERVL during MZT. **(D)** Expression level of MERVL copies in nontarget and writer KD samples in middle 2-cell and morula. **(E)** Decay rate of MERVL RNA at the 4-cell stage in nontarget and knockdown embryos tested by qPCR with Act D treatment. Error bars indicate mean +SD *, p-value<0.05, *** p-value<0.001. **(F)** MERVL RNA levels quantified by qPCR in the middle 2-cell and 4-cell embryos of control and dCas13b-ALKBH5 with gRNAs targeting MERVL. Error bars indicate mean ± SD. **(G)** Schematic diagram of m^6^A function in transcripts of genes and TEs during MZT. m^6^A+ maternal genes are downregulated when m^6^A writers are depleted in oocytes. M^6^A methylation promotes mRNA degradation of 2C genes after the 2-cell stage. Maternal MTA elements with m^6^A modification are highly expressed in oocytes. Kiaa1429 m^6^A writer knockout results in downregulation of MTA RNA without a change in chromatin accessibility. MERVL transcripts are highly modified by m^6^As, which control its RNA degradation after 2 cells.

In this study, we first developed an ultralow-input MeRIP-seq (ULI-MeRIP-seq) method, which enabled the profiling of m^6^A RNA methylation in mouse oocytes and preimplantation embryos. Our study not only generated the highly dynamic landscape of m^6^A on both coding and repetitive RNAs but also uncovered various roles of m^6^As during MZT: keeping maternal mRNAs stable and promoting decay of 2C-specific mRNAs (Fig. 4G). Importantly, our data on MTA and MERVL highlighted the timely storage and clearance of transposable elements through m^6^A-mediated posttranscriptional regulation, which may also play important roles in multiple developmental events.

## Acknowledgments

We thank Dr.C.He for the suggestion on the project. We thank Dr.R,Le for helpful advice on manuscript. Thank Dr,C.Ning. for the help with bioinformatic analysis. Thank Dr.H,Wang for providing the dCas13b-ALKBH5 plasmid.

## Author contributions

YG and SG designed project and directed all the experiments and bioinformatics analyses. BS provided advice and designed conditional knockout mice. YW carried out most experiments. XX carried out most bioinformatics analyses. MQ and ML carried out generation, breeding and genotyping of Kiaa1429 ^Zp3-cKO^ mice and Ythdf2^Zp3-cKO^ mice. YW and CC developed ULI-MeRIP-seq method. MQ and BS performed mini-ATAC on oocyte. RY and XL helped with embryo collection. WL, XK and YZ performed siRNA microinjection to embryos. MZ and CY performed LC-MS/MS for quantification of m^6^A level in embryos. HW helped with cell culture work. YG, YW, XX and SG wrote the manuscript with input from all authors.

## Data and materials availability

All data are available in the main text or the supplementary materials. All the MeRIP-seq, RNA-seq and ATAC-seq data generated in this study are summarized in Supplementary Table 1. The accession number for the sequencing data reported in this paper is GSA: CRA003985. These data have been deposited in the Genome Sequence Archive(*48, 49*) under project PRJCA004536. The shared URL for review is: https://bigd.big.ac.cn/gsa/s/v9YicRas.

## Supplementary Materials

Materials and Methods

Figs. S1 to S9

Tables S1 to S7

References (*36–49*)

