## Supplementary material for "The dual role of N6-methyladenosine on mouse maternal RNAs and 2-cell specific RNAs revealed by ULI-MeRIP sequencing": methods and supplemental figures

### Materials and Methods

*Ythdf2$^{Zp3}$* cKO mice generation

We purchased the targeted ES cell clone from EUCOMM to generate Ythdf2 KO first chimeric mice. Then, the chimeric mice were mated with Flpe mice to delete the Neo cassette to generate Ythdf2-floxed mice. We generated oocyte-specific knockout mice by crossing Ythdf2-floxed mice with Zp3-Cre mice. *Ythdf2$^{Zp3}$* cKO mice were generated and housed in Nanjing Medical University, Nanjing, China.

Animals and mouse embryo collection

Specific-pathogen-free (SPF) mice were housed in the animal facility at Tongji University, Shanghai, China. All animal maintenance and experimental procedures were performed according to Tongji University Guide for the use of laboratory animals. We collected GV oocytes from ovaries at 48 hrs after 5 IU of pregnant mare serum gonadotropin (PMSG) induction. Fully grown GV oocytes were collected in M2 medium with 2.5 μM milrinone after pricking the antral follicles. Cumulus cells around the oocytes and zona pellucida (ZP) were removed after 0.5% protease E digestion, and the oocytes were repeatedly drawn and blown in M2 medium (Sigma, M7167) with 2.5 μM milrinone using a mouth pipette. MII stage oocytes were collected from the oviducts of superovulated mice (with 5 IU of PMSG injection and 6 IU human chorionic gonadotropin (hCG) injection after 48 hrs. MII were collected 16 hrs after hCG injection). IVF proceeded immediately after MII oocyte collection and capacitation of sperm for 20 min in IVF-PLUS medium (Vitrolife, 10136). *In vitro* fertilization was conducted in IVF-PLUS medium for 4 hrs. Excessive sperm were removed after 4 hrs of IVF. Fertilized embryos (with a second polar body) were picked and cultured in CZB medium. All embryos and oocytes were digested in 0.5% protease E (Sigma, P8811) for ZP removal and washed in 0.5% BSA-PBS several times before being transferred into RNAiso plus reagent (Takara, 9109). Detailed information, including the background of embryos and the time of collection, is listed in table S6. *Kiaa1429$^{Zp3}$* cKO mice were from Bin Shen's lab(*21*).

When testing the RNA lifetime of specific RNAs, we transferred the 4-cell embryos (IVF 48 hrs) into Actinomycin D (Sigma, A9415) in CZB medium and cultured them for 6-8 hrs. The concentration of Actinomycin D for 4-cell embryo treatment was tested (fig. S7I). Finally, embryos were collected at 0 hrs, 3 hrs and 6 hrs with 1 μg/mL Actinomycin D treatment.

RNA extraction

ES cells or embryos were lysed in RNAiso plus reagent. A half volume of chloroform was extracted and transferred to a MaXtract tube (Qiagen, 129046). After 10 minutes of spin at full speed, the aqueous phase was recovered and transferred to a new tube. A 1/10 volume of 3 M NaAc and 2 μl glycogen (5 mg/mL, Roche, 10901393001) and an equal volume of isopropanol were added for RNA precipitation. The mixture was incubated at -80℃ for 30 minutes. RNA was precipitated after centrifugation for 15 min at full speed. After washing twice with 75% ethanol, the RNA pellet was resolved in RNase-free water.

Ultralow input MeRIP (ULI -MeRIP)

ULI-MeRIP was performed according to a previously described protocol (*24*) with some modifications. Briefly, 1 μg anti-m$^6$A antibody (Millipore, ABE572) was incubated with 10 μl Dynabeads protein A and 10 μl Dynabeads protein G (Invitrogen) and rotated for 2 hrs in IP buffer (150 mM NaCl, 10 mM Tris-HCl [pH 7.5], 0.1% NP40 in nuclease-free H2O) for one IP reaction. After incubation, beads were washed twice in IP buffer.

Before immunoprecipitation, RNA was fragmented by sonication. RNA (approximately 150 ng of embryo RNA) was extracted with 3 µl 1:1000 $m^6A$+ control RNA (GLuc), 3 µl 1:1000 $m^6A$- control RNA (CLuc), 150 ng carrier ssDNA (if low input RNA, sequence in table S7), 1 µl RNase inhibitor (40 U/µl), 1 µl SUPERase inhibitor (20 U/µl) and nuclease-free water to 50 µl and fragmented to 200-600 bp using Covaris S220. A 1/20 volume of fragmented RNA mixture was kept as input. The remaining sample was diluted in 600 µl IP buffer for 3 replicates of IP reactions. Each 200 µl dilution mixture was incubated with antibody-coated beads as one IP reaction and rotated at 4°C for 4-5 hrs.

After washing with IP buffer twice, low salt buffer (50 mM NaCl, 10 mM Tris-HCl [pH 7.5], 0.1% NP40 in nuclease-free $H_2O$) twice and high salt buffer (500 mM NaCl, 10 mM Tris-HCl [pH 7.5], 0.1% NP40 in nuclease-free $H_2O$) twice for 5 minutes each time. After washing, RNA-antibody-coated beads were resuspended in RNAiso plus reagent. RNA was extracted the same way as RNA extraction. Eluted RNA was used for RT-qPCR or library preparation.

Library preparation of ULI-MeRIP-seq and total RNA-seq

Library preparation of ULI-MeRIP samples and total RNA samples using SMARTer Stranded Total RNA-Seq Kit version 2 (Takara, 634411) was performed according to the manufacturer's protocol. Input RNA and IP extracted RNA were reverse transcribed to cDNA without fragmentation. Unfragmented samples were subjected to a fragmentation step according to the manufacturer's protocol. Libraries of IP and input samples (or total RNA samples) were amplified for 16-19 cycles and 14-15 cycles, respectively. After amplification, libraries were purified by AMPure XP beads. Purified libraries were sequenced on an Illumina NovaSeq 6000 platform (Novogene Corporation).

RT-qPCR

For ESC samples, cDNA was reverse-transcribed using an All in One Reverse Transcription Kit (ABM, G490) directly after ULI-MeRIP. For embryo samples, GLuc and CLuc expression was tested using reverse transcription products by qPCR. mRNA expression was tested using a diluted sequencing library. qPCR primers are listed in table S7.

Quantitative analysis of $m^6A$ level

For the quantification of $m^6A$ levels in embryos, 50 ng of extracted total RNA was injected into an LC-MS/MS instrument, which included ultra-performance liquid chromatography with a C18 column and triple-quadrupole mass spectrometry (AB SCIEX QTRAP 5500). The $m^6A$ level was detected in positive ion multiple reaction-monitoring (MRM) mode and quantified by nucleoside to base ion mass transitions (282.0 to 150.1 for $m^6A$ and 268.0 to 136.0 for A). The $m^6A$ concentration was calculated from the standard curve, which was generated from pure nucleoside standards.

miniATAC-seq

The ATAC-seq libraries were prepared as previously described with minor modifications[34]. Briefly, the nuclei of 30-50 GV oocytes were extracted by a Piezo-actuated micropipette and transferred to 10 µl lysis buffer (3.3 µl 1xPBS, 1.15 µl RNase-free H2O, 5 µl 2 x TD buffer, 0.25

µl Tn5 (Vazyme, TD501), 0.1 µl 1% Digitonin, 0.1 µl 10% Tween 20, 0.1 µl 10% NP40). Samples were mixed thoroughly by pipetting up and down several times and then incubated at 37°C at 1,000 rpm for 30 min. Then, 100 µl Proteinase K buffer (0.1 M Tris-HCl pH8.0, 0.2 M NaCl, 5 mM EDTA, 0.4% SDS) and 2 µl Proteinase K (Thermo Fisher Scientific, EO0491) were added to the samples and incubated at 55°C for 1 hour. To purify DNA, we added 50 ng of carrier RNA (Thermo Fisher Scientific, 4382878) to the samples and used phenol-chloroform to extract genomic DNA. Dr.GenTLE Precipitation Carrier (Takara, 9094) was used to precipitate DNA. To eliminate mitochondrial DNA, we added Cas9 protein and CARM sgRNA library[(35)] to the samples and incubated at 37℃ for 30 min, then amplified the libraries for 20-22 cycles. Finally, the libraries were purified using DNA Clean beads by 0.5×/1.5× double size selection.

dCas13b-ALKBH5 based *in vivo* $m^6A$ removal assay
The dCas13b-NES-ALKBH5 plasmid was a gift from Dr. Hongsheng Wang (Sun Yat-sen University). T7-promoter-dCas13b-NES-ALKBH5 was inserted into the pUC-GW vector by Genewiz company. *In vitro* transcription of dCas13b-NES-ALKBH5 was performed using the mMESSAGE mMACHINE Kit (Ambion, AM1340) after linearization of the template plasmid. gRNA was designed at $m^6A$ conserved regions according to the $m^6A$ MeRIP seq results. gRNA was synthesized by GenePharma company. Sequence of gRNA was listed in table S7. gRNA (20 µM each) and dCas13b-NES-ALKBH5 mRNA (200 ng/µl) were mixed and injected (approximately 10 pl) into oocytes by a Piezo-driven micromanipulator. *In vitro* fertilization was performed immediately after microinjection. Embryos are collected at 2-cell stage. $m^6A$ MeRIP qPCR was performed to validate the efficiency of $m^6A$ removal by MERVL RNAs.

Knockdown of Mettl3/14/16 in early embryos
The nontarget mixture contained igG antibody (200 ng/µL, Millipore, 12-371) and siNontarget (5 µM). The knockdown mixture contained anti-METTL3 antibody (200 ng/µL, Abcam, ab195352), anti-METTL14 antibody (200 ng/µL, Sigma, E115616), anti-METTL16 antibody (200 ng/µL, Abcam, ab186012), siMettl3 (5 µM), siMettl14 (5 µM) and siMettl16 (5 µM each). The mixture was prepared and centrifuged for 10 min. The siRNA sequences are listed in table S7 and synthesized by GenePharma company. MII oocytes from BDF1 female mice (C57BL/6 female mice cross DBA/2 male mice) were injected with approximately 10 pL of mixture solution using a Piezo-driven micromanipulator. IVF proceeded after 2 hours of injection. Fertilized embryos were moved to CZB medium for culturing.

Single-cell RNA-seq.
The process of single-cell RNA-seq for *Ythdf2* $^{Zp3}$ cKO embryos was performed as previously described(*36, 37*). Briefly, *Ythdf2* fl/fl (Ctrl) and fl/fl; Zp3-cre (cKO) oocytes and 2-cell embryos were harvested and washed three times in 0.5% BSA/PBS. Then, oocytes or embryos were transferred into 4 µl of lysis buffer (1x PCR Buffer II, 3 mM MgCl2, 0.45% NP40, 4.5 mM DTT, 0.36 U/µl RNase inhibitor, 0.18 U/µl SUPERase inhibitor) using a mouth pipette. Primer (0.5 µM V1-T24) and dNTPs (2.5 µM) were added to the lysate after lysis at 70°C for 90 seconds. Reverse transcription was performed directly using SuperScript III. Unreacted primers were digested using Exo-SAP-IT. Terminal deoxynucleotidyl transferase (TdT) was then used to add a poly(A) tail onto the 3' end of the first-strand cDNAs. The total cDNA library was then amplified by PCR (16-19 cycles). The amplified cDNA library was fragmented using the Covaris S220 sonicator. The sequence libraries were prepared using a KAPA HyperPlus Library

Preparation kit (Roche, kk8504). Single-end 50-bp sequencing was performed on an Illumina HiSeq 2500 at Berry Genomics Corporation.

RNA-seq data processing

RNA-seq data were first subjected to Trim_galore (version 0.6.4) for adaptor trimming as well as quality control with the parameters --paired -j 7 --basename. The trimmed paired-end reads were then aligned to the mm9 reference genome with random chromosomes cleaned by Hisat2(*38*) (version 2.1.0) under the parameters -p 15 --dta-cufflinks --no-mixed –no-discordant. The expression of genes was quantified as FPKM by Cufflinks(*39*) (version 2.2.1). For the downstream data analyses, FPKM values were averaged for each gene between replicates. The RefSeq gene annotation files were downloaded from UCSC. For genes with multiple isoforms, the longest transcripts were selected. The R package DESeq2(*40*) (version 1.26.0) was used for gene differential expression analysis. Fold change > 2 and FDR < 0.05 were used as cutoffs for downregulated and upregulated genes. Genome coverage bigwig files for the UCSC genome browser were generated by deeptools(*41*) (version 3.5.0) bamCoverage with parameters –normalizeUsing CPM -bs 1. Genome coverage bigwig files for aggregation plots were generated by bamCoverage with parameters –normalizeUsing RPKM -bs 50. An aggregation plot for the input signal was plotted by computeMatrix and plotProfile functions of the Deeptools package.

ULI-MeRIP data processing

The strategy for sequencing read trimming and mapping of ULI-MeRIP data was the same as that of RNA-seq. The $m^6A$ peaks in each $m^6A$ immunoprecipitation sample were identified by the MACS2(*42*) (version 2.2.4) callpeak tool with the corresponding input sample serving as a control. MACS2 was run with options '-g mm --nomodel --keep-dup all' for each replicate. For downstream analysis, a peak was kept if it was shared in all replicates. $m^6A$-modified genes were defined as if any exon of that gene overlapped with a $m^6A$ peak. Genome coverage bigwig files for the UCSC genome browser were generated by deeptools bamCoverage with parameters –normalizeUsing CPM -bs 1. Genome coverage bigwig files for aggregation plots were generated by bamCoverage with parameters –normalizeUsing RPKM -bs 50. Aggregation plot for the input signal was plotted by computeMatrix and plotProfile functions of the Deeptools package.

Consensus motif identification within $m^6A$ peaks

High confidence peaks with fold change greater than 4 were used to search the enriched motif for genes or repeats. The coordinates of peak-summit regions (50 nt upstream and downstream flanking the summit) of high confidence peaks were retrieved, and those peak-summit regions were mapped to the annotated genes or repeats to fetch the strand information by bedtools. Then, the peak-summit regions with strand information were subjected to findMotifsGenome.pl of the HOMER(*43*) suite to identify motifs under parameters -rna -len 6.

Peak enrichment analysis

Enrichment of $m^6A$ peaks in gene-related genomic elements and repetitive elements was calculated as

log2 ratio of the observed over expected counts of $m^6A$ peaks overlapping with selected genomic regions. The observed counts were calculated as the number of $m^6A$ peaks overlapping with related genomic regions using intersectBed of bedtools by restricting that the proportion of overlapped region is greater than 50%. The expected counts were calculated as the number of random peaks generated using shuffleBed overlapping with related genomic regions. The radar plot for peak enrichment was plotted by the R package ggradar.

Gene group categorization

Maternal decay genes were defined as FPKM in GV or MII stage >5 and fold change (GV/2-cell) >4. ZGA genes were defined as FPKM in GV and MII <1, FPKM in zygote and 2-cell >1, and fold change (2-cell/GV) >4. Maternal decay genes and ZGA genes were further grouped according to their m$^6$A modification status. Genes meeting the criteria of both input FPKM >1 and overlapping with an exon m$^6$A peak were defined as m$^6$A-modified genes. A maternal decay gene was defined as m$^6$A+ if this gene was modified in the GV or MII stage; otherwise, it was defined as an m$^6$A- gene. Similarly, a ZGA gene was defined as m$^6$A+ if this gene was modified in the zygote or 2-cell stage; otherwise, it was defined as an m$^6$A- gene.

Gene ontology analysis

Gene ontology (GO) analysis of genes was performed using the compareCluster function of the R package clusterProfiler(*44*) (version 3.14.3) under the parameters fun="enrichGO", pAdjustMethod = "BH", ont= "BP", OrgDb= org.Mm.eg.db, keyType= 'SYMBOL', and qvalueCutoff = 0.05. Enrichment maps were constructed using the R package Complexheatmap (version 2.2.0). GO terms for each functional cluster were summarized to a representative term.

ULI-MeRIP-seq analysis of *Kiaa1429*$^{Zp3\text{-}cKO}$ GV oocytes

To determine the m$^6$A level after *Kiaa1429* knockout, the whole m$^6$A level was calculated by the number of reads mapped to the mouse genome divided by the number of reads mapped to m$^6$A-modified spike-in (GLuc). To normalize the input and IP signal after *Kiaa1429* knockout, the genome coverage bigwig files of KO IP and input were generated by bamCoverage with parameters –normalizeUsing RPKM -bs 50 –scaleFactor f. The scale factor f is calculated as:

$$f=(N^{WT}*T^{KO})/(T^{WT}*N^{KO})$$

T represents the library sequence depth, and N represents the read number of spike-ins. For input, N is the number of reads mapping to CLuc. For IP, N is the number of reads mapping to GLuc.

The first step of peak calling was the same as previously described, and peaks were further filtered by calculating the IP/input signal ratio by using scaled genome coverage bigwig files. Peaks with IP/input ratios greater than 1.5 were kept for further analysis.

Transposable elements analysis

Repeat elements were mm9 version of RepeatMasker annotation downloaded directly from UCSC (http://hgdownload.cse.ucsc.edu/goldenPath/mm9/database/rmsk.txt.gz). Alignment was performed using hisat2 against mm9 under the parameters -p 15 --dta-cufflinks --no-mixed –no-discordant. For multiple mapping, up to 5 alignments for each read were reported. We excluded repeats that overlapped with genic regions to rule out the impact of gene expression. To quantify the expression level for each locus, the reads per kilobase per million mapped reads (RPKM) value for each genome-wide nonoverlapping 50 bp window was calculated as the signal density in sequencing data by the bamCoverage program with parameters –normalizeUsing RPPM -bs 50, and RPKM values were averaged for windows covering the locus and further averaged between replicates. The expression level of each family was the average level of all copies. For

downstream analysis, families with expression levels greater than 5 in at least one stage were kept.
To plot the aggregation profiles, full-length MTA and MERVL annotation were applied. Full-length MTA was defined as MTA_Mm, MTA_Mm-int, and MTA_Mm from a previously published paper(*45*) using mergeBed, allowing at most a 20 bp gap between adjacent annotations. Full-length MERVL was defined as MT2_Mm, MERVL-int, ORR1A3-int, MERVL-int, and MT2_Mm from a previously published paper(*46*) using the same method as MTA.

ATAC-seq data analysis
ATAC-seq data were aligned by bowtie2 (version 2.3.5.1)(*47*) by the parameter --no-mixed --no-discordant. PCR duplicates were removed by Picard (version 2.21.1) function MarkDuplicates with default parameters. MTA elements were divided into 5 groups according to RNA expression level (0-1, 1-5, 5-10, 10-50, > 50). The ATAC-seq signal of each group was measured by the average RPKM. One MTA element (chr2:98509266-98510936) contained an unusual tall, short peak of read density and was excluded from analysis. Promoters were defined as TSS +-2 kb. To calculate ATAC signals at promoters, genome coverage bigwig files for aggregation plots were generated by bamCoverage with parameters –normalizeUsing RPKM -bs 50, RPKM values were averaged for windows covering the locus and further averaged between replicates.

Statistical analysis
R (version 3.6.1) was used for statistical analysis. Pearson's correlation coefficient was calculated using the cor.test function with default parameters to evaluate the reproducibility of replicates. The Wilcoxon rank sum test was used for group comparisons using the wilcoxon test function in R. * represents $p < 0.05$, ** represents $p < 0.01$, and *** represents $p < 0.001$. Fisher's exact test was used for GO enrichment analysis by clusterProfiler. The number of embryos or biological replicates used in this study can be found in table S6 and the Methods Details section. No statistical methods were used to predetermine the sample size.

Code availability
All analyses were made based on Shell and R codes and are available upon request.

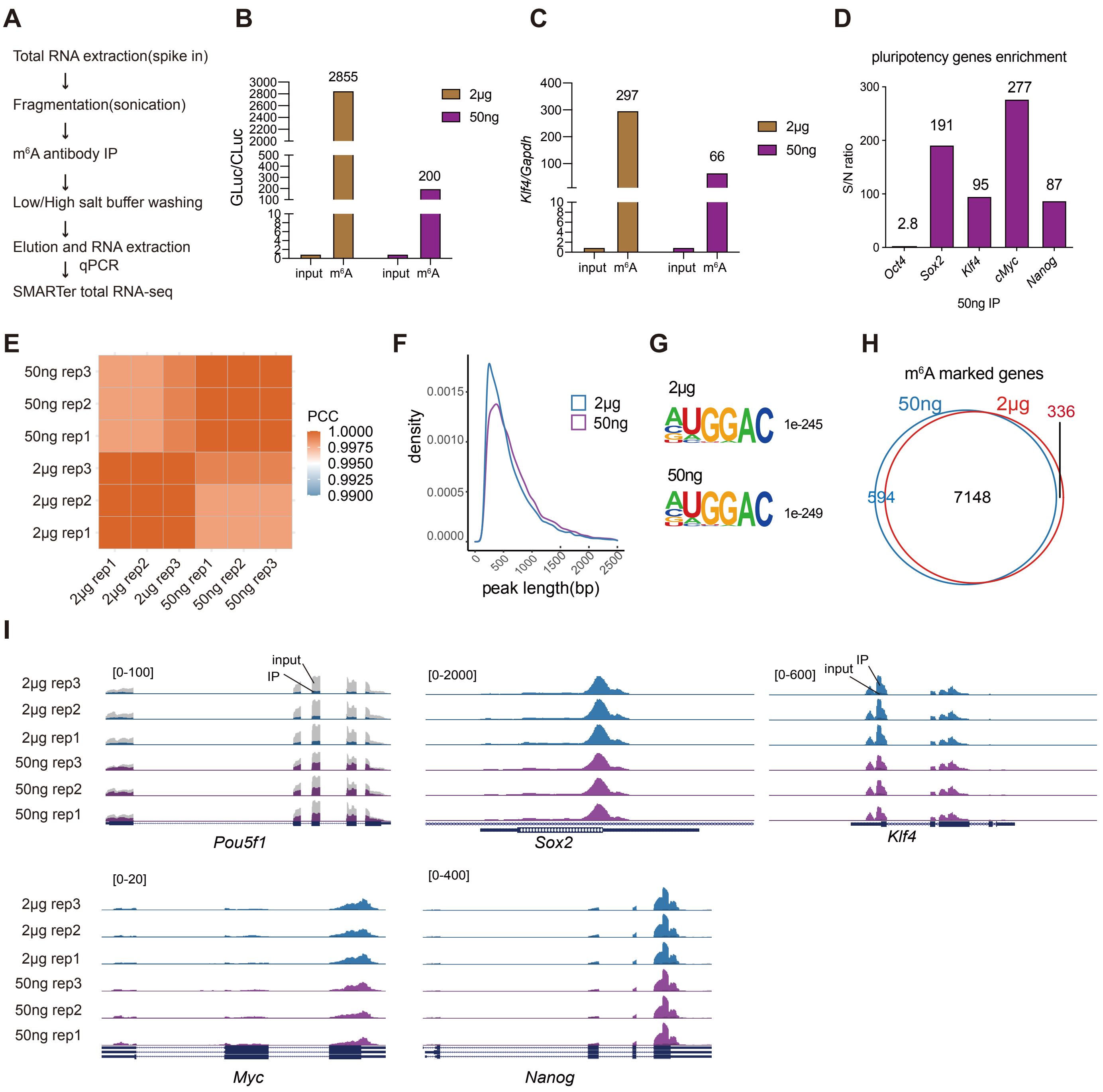


**fig. S1. Validation of ULI-MeRIP-seq in mESC.**

(**A**) Workflow of ULI-MeRIP-seq. (**B** and **C**) High enrichment of m$^6$A in IP samples with 2 μg and 50 ng total RNA tested by qPCR of GLuc vs CLuc (B) and *Klf4* vs *Gapdh* (C). (**D**) S/N ratio[24] of pluripotency genes in 50 ng total RNA of mESCs tested by qPCR. S/N, signal-to-noise ratio, represents IP over input of target genes vs *Gapdh*. (**E**) The Pearson correlation coefficients (PCCs) showing a high correlation of the m$^6$A signal from ULI-MeRIP-seq data between 50 ng and 2 μg starting amount of total RNA in mESCs. (**F**) Density of m$^6$A peak length in 2 μg and 50 ng starting RNA amount. (**G**) Sequence logo and p-values of the consensus motif of m$^6$A peak centers located at genes. (**H**) Venn plot showing the overlapping m$^6$A-marked genes between 2 μg and 50 ng starting RNA amounts in mESCs. (**I**) UCSC browser tracks showing IP and input read distribution of representative pluripotency genes in 2 μg and 50 ng samples.

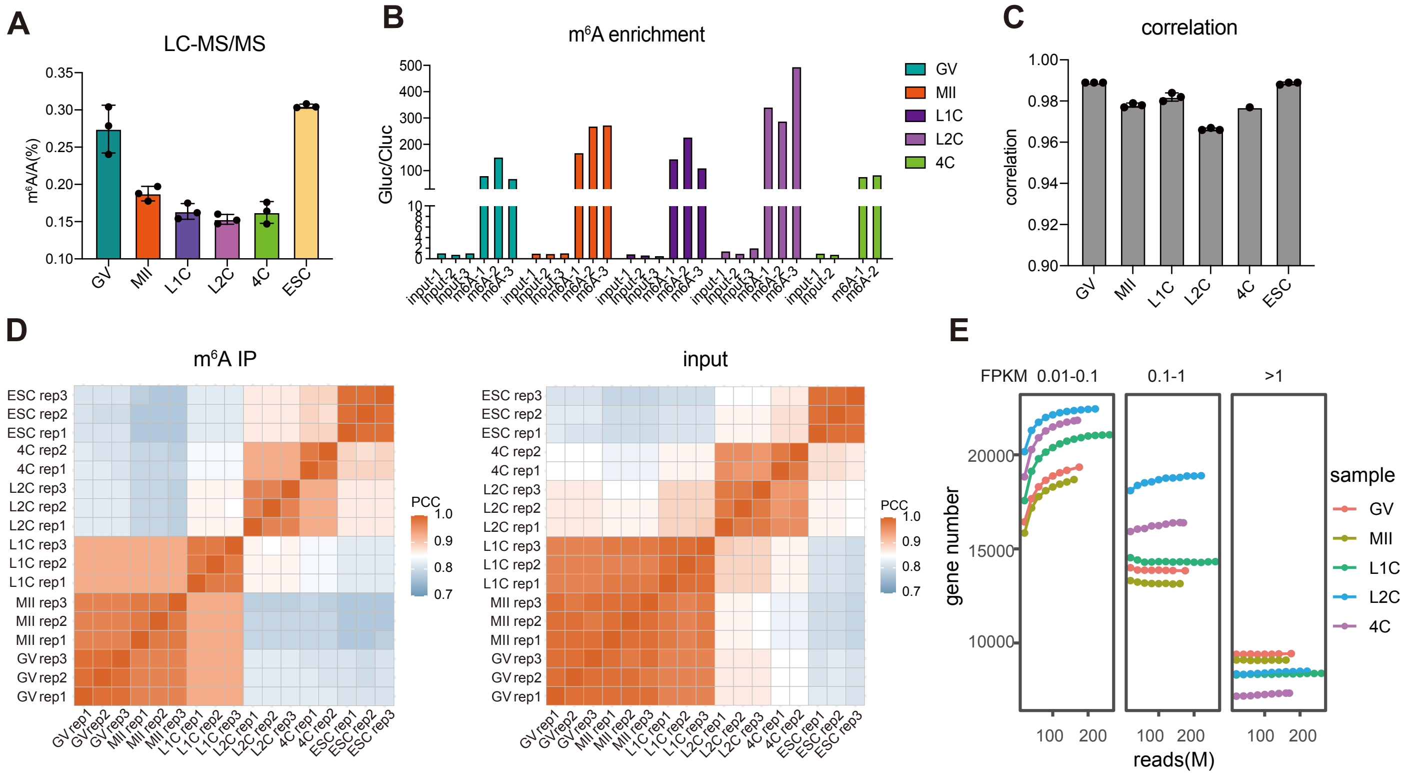


**fig. S2. Validation of ULI-MeRIP-seq data quality in oocytes and embryos.**

(**A**) Low-input LC-MS/MS quantification of the $m^6A/A$ ratio in total RNAs (approximately 50 ng), n = 3 biological replicates. Error bars indicate mean ± s.e.m. (**B**) High enrichment of $m^6A$ in embryo IP samples tested by qPCR of GLuc vs CLuc. (**C**) PCC of two or three replicates in IP samples. Error bars indicate mean ± s.e.m. (**D**) Heatmap showing PCC of IP and input samples. (**E**) Saturation plot showing that sequencing reads can cover high (FPKM>1), medium (0.1<FPKM<1) and low (0.01<FPKM<1) levels of genes in input.

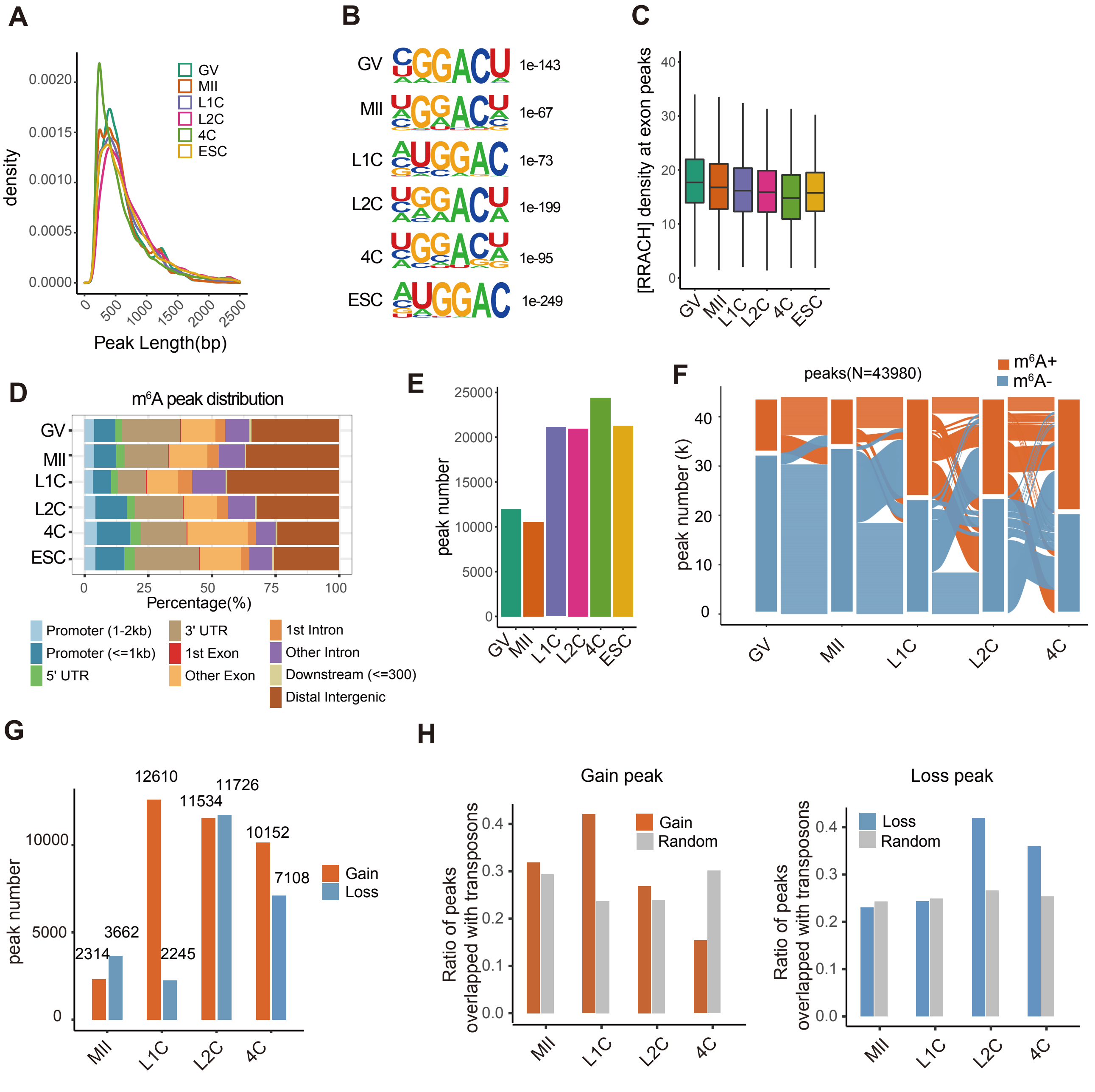


**fig. S3. Features of RNA m⁶A modification in mouse oocytes and embryos.**

(**A**) Density of m⁶A peak length in oocytes, early embryos and mESCs. (**B**) Sequence logo and p-values of the consensus motif of m⁶A peak centers in each sample. (**C**) [RRACH] motif density of peaks at exons in each sample. (**D**) Bar chart presenting the fraction of m⁶A peaks in different genomic regions. (**E**) Bar chart showing m⁶A peak number in each sample. (**F**) Alluvial diagram showing the global dynamics of m⁶A peaks during MZT (N=43980). Each line represents a m⁶A peak in at least one stage. (**G**) The number of peaks gaining or losing m⁶A at each stage compared to the prior stage during MZT. The number of peaks gaining or losing m⁶A is indicated above each bar. (**H**) Bar plot showing the ratio of gained or loss peaks overlapping with transposable elements compared to random peaks.

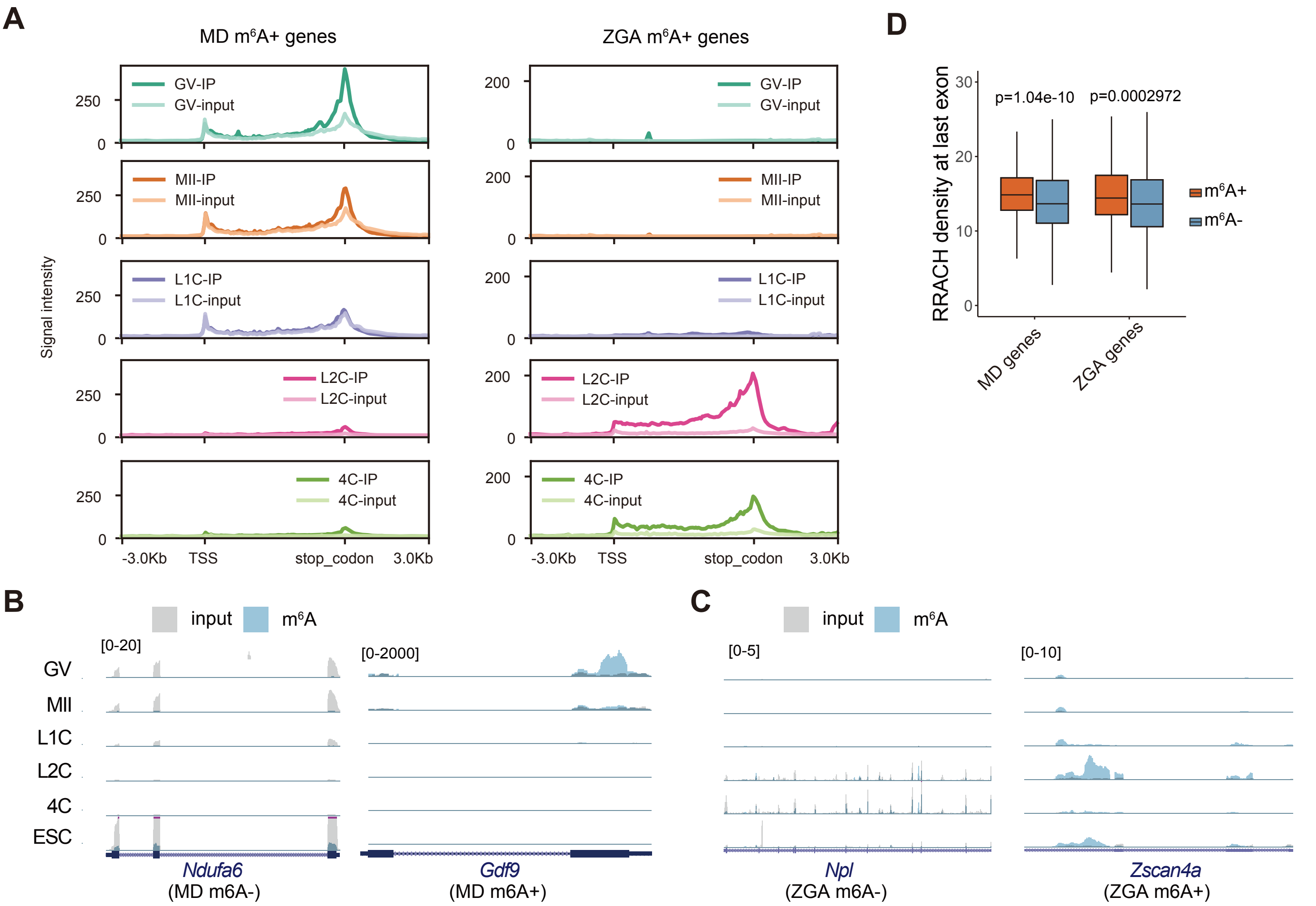


**fig. S4. Properties of MD m⁶A+ genes and ZGA m⁶A+ genes.**

(**A**) Average profile of m⁶A IP and input signal of maternal decay (MD) m⁶A+ genes and ZGA m⁶A+ genes during MZT. (**B** and **C**) The UCSC browser track showing IP and input reads of example genes. (**D**) Box plot showing [RRACH] motif density at the last exon in MD genes and ZGA genes with and without m⁶As.

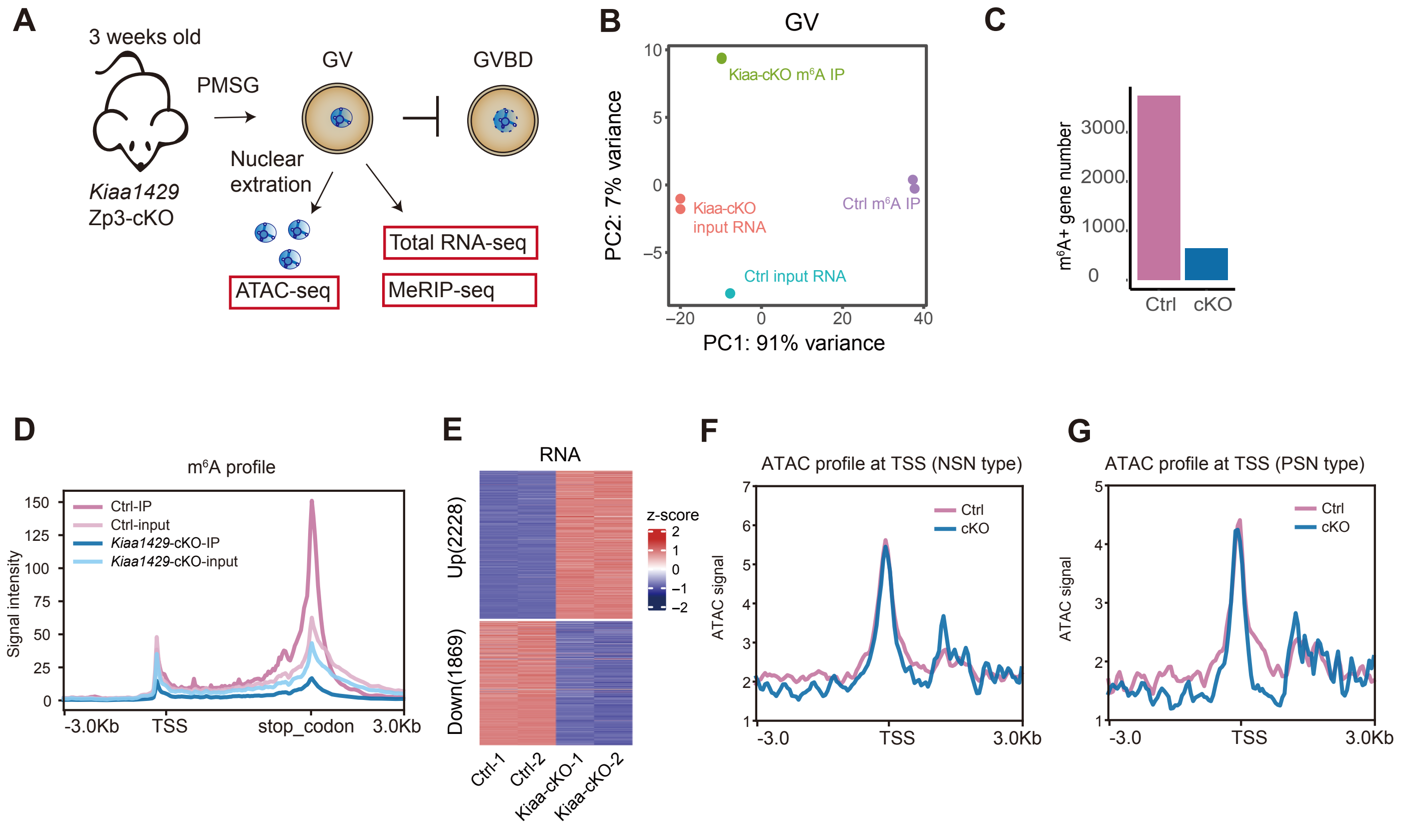


**fig. S5. m⁶A maintains the high expression levels of marked genes during oocyte development.**

(**A**) Schematic diagram of total RNA-seq, m⁶A MeRIP-seq and nuclear ATAC-seq using *Kiaa1429* Zp3-cKO GV oocytes. *Kiaa1429* Zp3-cKO GV oocytes arrest before germinal vesicle breakdown (GVBD).

(**B**) PCA of m⁶A IP and input of control and *Kiaa1429* Zp3-cKO GV oocytes.

(**C**) Bar chart showing m⁶A+ gene number in control and *Kiaa1429* Zp3-cKO GV oocytes.

(**D**) Average m⁶A IP and input signal in control and *Kiaa1429* Zp3-cKO GV oocytes at m⁶A+ genes of WT GV. The IP and input signal were scaled by GLuc and CLuc spike-ins, respectively.

(**E**) Heatmap showing the normalized expression level of DEGs between *Kiaa1429* Zp3-cKO and control GV oocytes.

(**F** and **G**) ATAC signal at promoter regions (TSS -3 kb to 3 kb) of m⁶A+ MD genes in NSN (left) and PSN (right) type GV oocytes of control and *Kiaa1429* cKO mice.

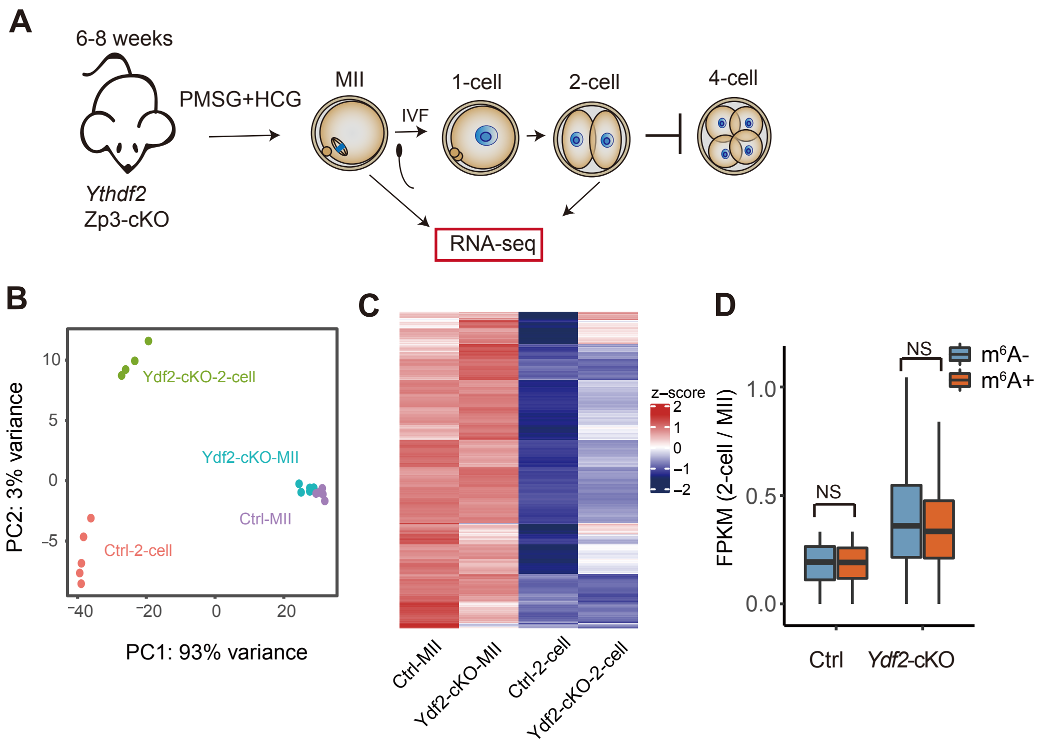


**fig. S6. *Ythdf2*$^{Zp3}$ cKO impairs maternal gene decay independent of m$^6$A modification.**

(**A**) Schematic diagram of RNA-seq for early embryos using *Ythdf2* Zp3-cKO mice. (**B**) PCA of RNA-seq for MII oocytes and 2-cell embryos of *Ythdf2* cKO and control mice. (**C**) Heatmap of normalized expression levels of maternal decay genes for MII oocytes and 2-cell embryos in control and *Ythdf2* cKO mice. (**D**) 2-cell over MII ratio of expression level (FPKM) for m$^6$A+ and m$^6$A- maternal decay genes in *Ythdf2* control and cKO mice. NS, p-value>0.05.

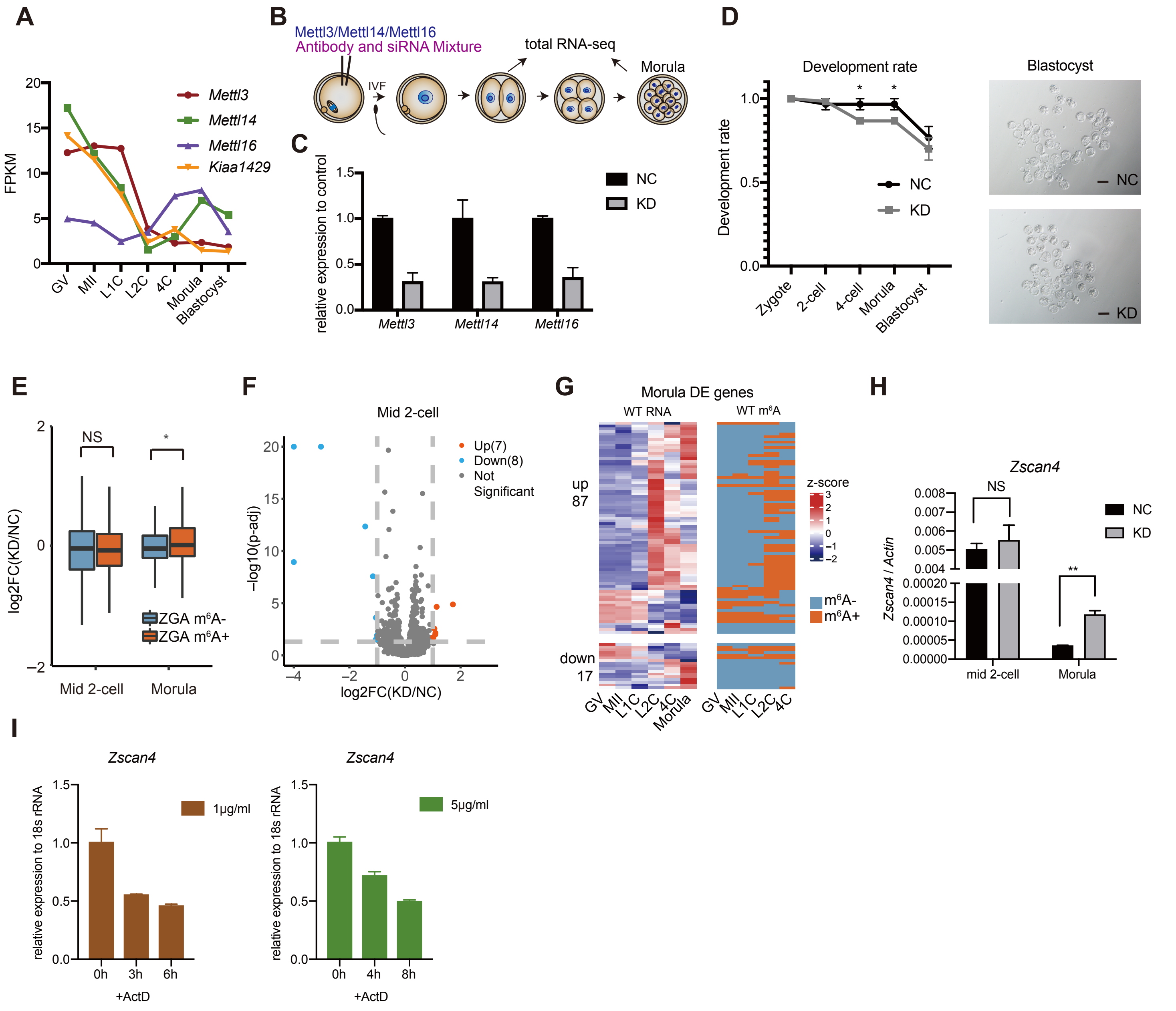


**fig. S7. Knockdown of m$^6$A writers in early embryos affects the decay of 2C genes.**

(**A**) Expression pattern of known m$^6$A writers *Mettl3*, *Mettl14*, *Mettl16* and *Kiaa1429* during early embryonic development. (**B**) Schematic diagram of writer knockdown (KD) assay. MII oocytes were injected with antibodies and siRNA mixture and then subjected to *in vivo* fertilization (IVF). Total RNA-seq was conducted for 2-cell and morula embryos. (**C**) Knockdown efficiency tested at morula stages by qPCR. NC, nontarget, injection with igG antibody and nontarget siRNA. KD, knockdown, injection with METTL3, METTL14 and METTL16 antibody and siRNA mixture. Error bars indicate mean ± SD. (**D**) Development rate (left) and blastocyst photos (right) in the NC and KD groups. Error bars indicate mean ± s.e.m. *, p-value<0.05. scare bar = 100μm. (**E**) Log2-fold change of KD over NC of ZGA m$^6$A+ and m$^6$A- genes in middle 2-cell and morula embryos. NS, p-value>0.05, *, p-value<0.05 (**F**) Volcano plots showing DEGs between writers KD and NC group in middle 2-cell. (**G**) Heatmap showing normalized expression level of DEGs between writers KD and NC group (left panel), and corresponding m$^6$A modification status in WT oocytes and embryos (right panel). (**H**) Expression level of Zscan4 tested by qPCR after knockdown of m$^6$A writers. NS, p-value >0.05, **, p-value < 0.01. Error bars indicate the mean ± s.e.m. (**I**) Expression level of Zscan4 relative to 18S rRNA after Actinomycin D treatment at different concentrations. Error bars indicate mean ± SD.

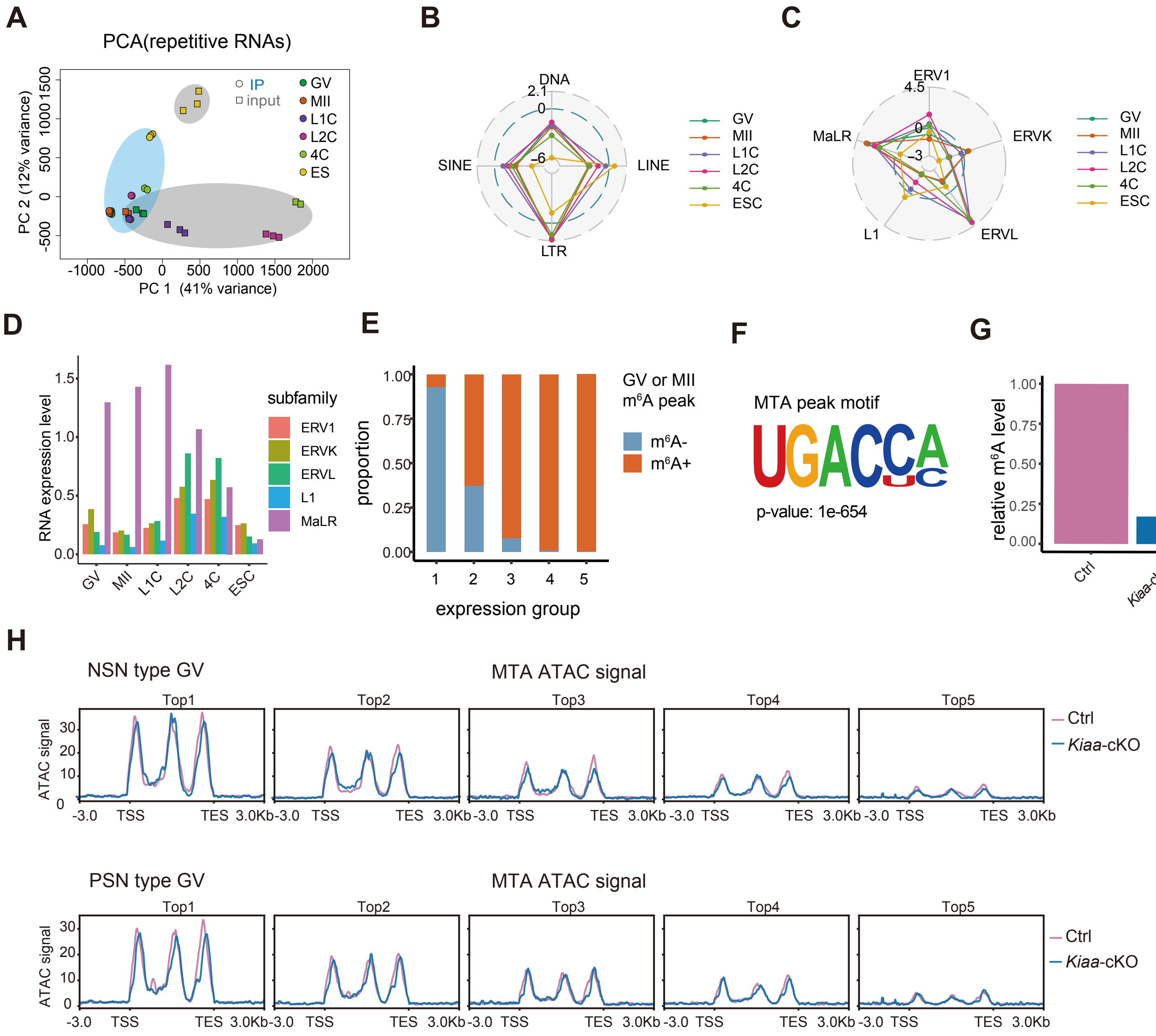


**fig. S8. m⁶A profiles of TEs and maternal-specific TE MTA**

(**A**) PCA showing m⁶A IP and input samples at all repetitive RNAs during MZT. Squares are input samples, and circles are m⁶A samples. (**B**) Radar chart showing the enrichment score (log2) of m⁶A peaks in four classes of transposable elements, including DNA transposons (DNA), SINEs, LINEs and LTRs. (**C**) m⁶A enrichment of major families of LINE and LTR shown by radar plot. (**D**) Average RNA level of ERV, L1 and MaLR during MZT. (**E**) Unmodified MTA copies are not expressed or of low expression level. Full-length MTA copies were classified into five groups according to expression levels. "1" represents copies with the lowest RNA level, while "5" represents copies with the highest level. Each group of copies was further classified according to m⁶A status. (**F**) Sequence logo and p-values of the consensus motif of m⁶A peak centers at the MTA locus. (**G**) Relative m⁶A signal intensity at MTA in GV oocytes of control and *Kiaa1429*-cKO mice normalized by GLuc. (**H**) ATAC signal of MTA in nonsurrounded nucleolus (NSN) and partly surrounded nucleolus (PSN) types of GV for control and *Kiaa1429*-cKO mice according to expression group. Top1 to Top5 represent five expression groups according to the expression level of MTA copies from high to low.

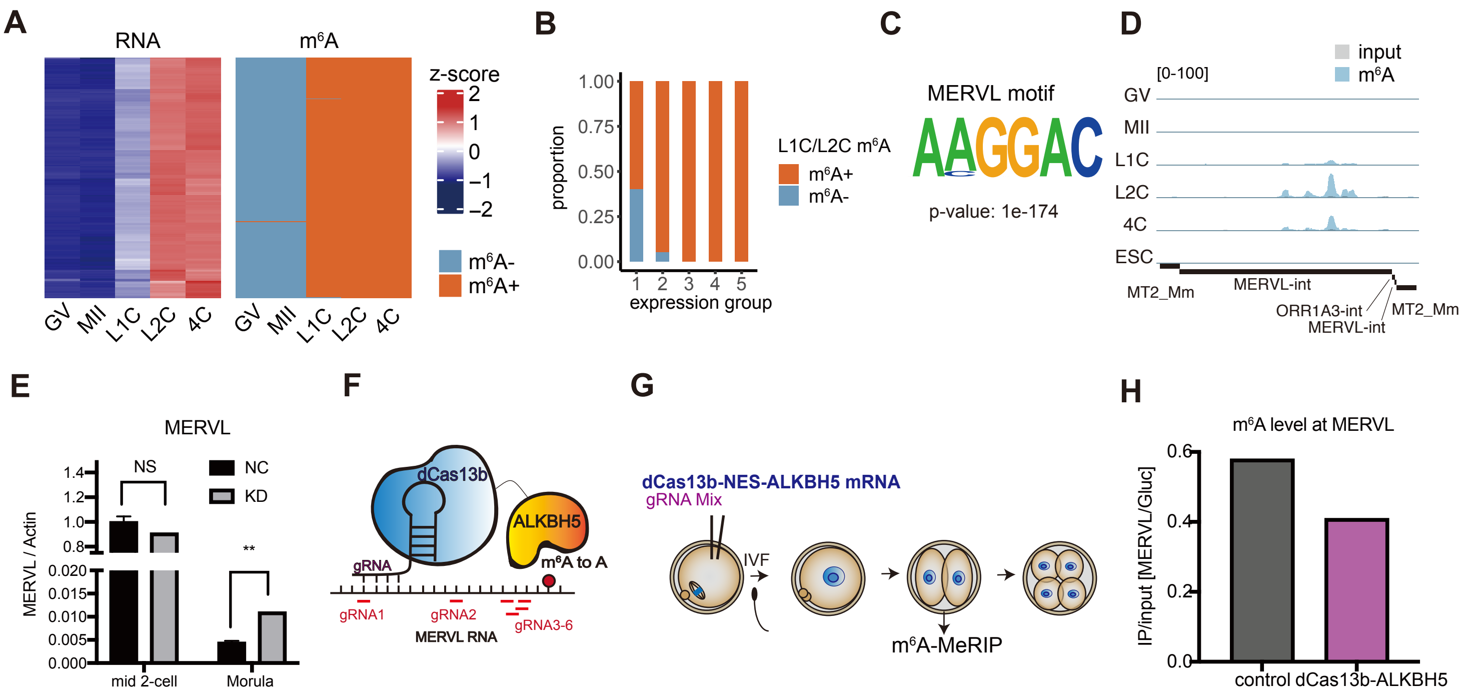


**fig. S9. m⁶A profiles of ZGA-specific TE MERVL.**

(**A**) Normalized RNA level (left panel) and m⁶A peak status (right panel) of full-length MERVL copies. (**B**) Full-length MERVL copies were classified into five groups according to expression levels. "1" represents copies with the lowest RNA level, while "5" represents copies with the highest level. Each group of copies was further classified according to m⁶A status. (**C**) Sequence logo and p-values of the consensus motif of m⁶A peak centers at the MERVL locus. (**D**) The UCSC browser track showing IP and input reads of MERVL during MZT. (**E**) Relative expression level of MERVL in nontarget and writer KD samples in middle 2-cell and morula quantified by qPCR. Error bars indicate mean ± s.e.m. NS, p-value>0.05, **, p-value<0.01. (**F**) Schematic model of the dCas13b-ALKBH5 site-specific demethylation system. Six gRNAs were used to target different sites of MERVL. (**G**) Schematic diagram showing the experimental design of the dCas13b-ALKBH5 MERVL demethylation assay. (**H**) The m⁶A level of MERVL was decreased after dCas13b-ALKBH5 targeting MERVL, as tested by m⁶A-MeRIP-qPCR.

Table S1. The mapping information for RNA and ATAC-seq samples.
Table S2. GLuc and CLuc reads number.
Table S3. Gene list for MD and ZGA genes.
Table S4. DEGs in Kiaa1429 Zp3-cKO GV oocytes.
Table S5. DEGs in writer knockdown 2-cell and morula.
Table S6. Embryo background and collection information.
Table S7. Oligo and primer sequence.
